# Metformin acts directly in the brain to slow features of neurodegeneration

**DOI:** 10.1101/2023.07.18.549549

**Authors:** Edward C. Harding, Hsiao-Jou Cortina Chen, Dmytro Shepilov, Marc Welzer, Viviana Macarelli, Alix Schwiening, Christine Rowley, Sanya Aggarwal, Dean Swinden, Deborah Kronenberg-Versteeg, Florian T. Merkle

## Abstract

Dementia is a largely untreatable syndrome that is epidemiologically associated with metabolic diseases such as type 2 diabetes (T2D) and obesity. Drugs used to treat T2D such as metformin are inexpensive, safely given to millions of people, and have also been reported to slow neurodegeneration. We hypothesised that the neuroprotective benefits of metformin might extend to metabolically healthy individuals and tested this hypothesis in a mouse prion model that recapitulates key features of human neurodegenerative disease, including synaptic loss and motor impairment. These features and the time course of this model (24 weeks) allows the effects of metabolic risk factors and metformin to be tested and potentially generalised to other forms of neurodegenerative disease. Mice fed a high fat diet (HFD) developed high adiposity with impaired glucose and insulin homeostasis, similar to the effects of chronic obesity seen in humans whereas mice on matched control diet (CD) remain metabolically healthy. Chronic treatment with metformin in HFD-fed mice significantly increased survival and health span relative to vehicle-treated mice. Mice fed a HFD also had a modestly extended health span relative to mice fed CD, as measured by development of motor signs of prion disease. Metformin also significantly extended health span in metabolically healthy CD-fed mice. Using targeted mass spectrometry, we found that metformin reached deep brain structures at functional concentrations, driving a reduction in pPERK and changing the activity of microglia *in vivo*. Metformin was able to alter ER stress pathways at the same concentrations in healthy animals and using human iPSC-derived microglia and mouse organotypic slices we show that the action of metformin at these concentrations does not require a systemic mechanism, necessary for the treatment of diabetes, and is likely a result of direct secondary pharmacology in the brain. Together, these data broadly support the premise of repurposing metformin for neuroprotection, even in metabolically healthy individuals.

## INTRODUCTION

Dementia is a devastating condition that remains largely untreatable. An ageing population, combined with health risk factors have accelerated the global prevalence of dementia (Brookmeyer *et al*., 2007) which now affects almost 70 million people (Nichols *et al*., 2022; Shin, 2022). Health risk factors for dementia include mid-life obesity (Ott *et al*., 1999; Kivimäki *et al*., 2018; Floud *et al*., 2020), which is predicted to affect nearly half of the US adult population by 2030 (Ward *et al*., 2019) and which also significantly increases the risk of cardiovascular disease and type 2 diabetes (T2D) (Algoblan, Alalfi and Khan, 2014). T2D affects more than 15% of the United States population (Wang *et al*., 2021) and is associated with a significantly increased risk of all-cause dementia (Ott *et al*., 1999), including Alzheimer’s Disease (AD) (Peila, Rodriguez and Launer, 2002; Arvanitakis *et al*., 2004; Crane *et al*., 2013), vascular dementia, and general cognitive decline (Ohara *et al*., 2011; Cheng *et al*., 2012).

In the brain, AD is characterised by the aggregation of misfolded tau protein into tangles, and the aggregation of β-amyloid protein into plaques (Murphy and LeVine, 2010). AD patients with diabetes have a greater β-amyloid plaque burden (Valente *et al*., 2010). The misfolding and aggregation of α-synuclein and prion protein are also key pathological features of Parkinson’s Disease (PD) and prion disease, respectively, suggesting that protein aggregation is a shared feature of common neurodegenerative diseases and may play a mechanistic role in disease progression. A variety of genetic and environmental factors contribute to neurodegenerative diseases but result in a common set of features including synaptic loss, endoplasmic reticulum (ER) stress, and neuron loss (Freeman and Mallucci, 2016; Wirths and Zampar, 2020). The association of metabolic and neurodegenerative disease seen in humans has been corroborated in animal models. Mice with chronic obesity or diabetes have cognitive deficits including memory impairments and the reduced expression synaptic markers such as PSD-95 relative to metabolically healthy controls (Kanoski and Davidson, 2010; Arnold *et al*., 2014; Gladding *et al*., 2018; McLean *et al*., 2018). Both mouse models of AD and post-mortem tissue from AD patients show consistent activation of insulin signalling components such as AKT serine threonine kinase 1 (AKT1) and hippocampal insulin receptor substrate 1 (IRS1), that correlate with the density of tau tangles and β-amyloid plaques (Arvanitakis *et al*., 2020).

Conversely and encouragingly, observation studies and patient clinical records suggest that the effective mitigation of risk factors, including diabetes and obesity, may reduce long-term dementia and could form new treatment practices to manage long-term dementia risk at a population level (Whitmer *et al*., 2008; Campbell *et al*., 2018; Samaras *et al*., 2020; Nørgaard *et al*., 2022; Heneka, Fink and Doblhammer, 2015; Tang *et al*., 2022). Specifically, drugs used to manage obesity and T2D have neuroprotective properties, though the underlying mechanisms are unclear. These drugs include metformin as well as agonists of the glucagon-like peptide 1 receptor (GLP1R) such as exenatide, liraglutide, and semaglutide which have shown neuroprotective effects in many animal studies (DiTacchio, Heinemann and Dziewczapolski, 2015; Yun *et* al., 2018; Yan et al., 2019; Zhang et al., 2019), and are in numerous human clinical trials such as ELAD (NCT01843075) (Femminella *et al*., 2019; Reich and Hölscher, 2022; Mantik *et al*., 2023). While these results are encouraging and support the concept that improving metabolic health can be neuroprotective, the long-term consequences of GLP1R agonists on cognition and brain health are still poorly understood. In contrast, the small molecule drug metformin has been a front-line diabetes treatment for decades and is one of the most widely prescribed drugs in the world. Furthermore, metformin has been associated with reduced cognitive decline in humans in both observational data (Samaras et al., 2020), and in randomised trials (Koenig *et al*., 2017; Mantik *et al*., 2023), with further trials ongoing including MetMemory (NCT04511416) and MET-FINGER (NCT05109169). Overall, metformin has considerable potential for repurposing in dementia supported by human, rodent and computational data (Shi *et al*., 2019; Zhang *et al*., 2021; Charpignon *et al*., 2022).

Here, we hypothesised that metabolic disease would accelerate disease progression in a mouse model of neurodegeneration, and that metformin would slow disease progression in both metabolically healthy and unhealthy mice. We tested this hypothesis in a mouse model of prion disease - Rocky Mountain Laboratory scrapie (RML) - that is characterised by a stereotyped progression that mimics human disease processes and features such as synaptic loss and cell death. We combined RML scrapie inoculation with long-term administration of a high-fat diet (HFD) to induce neurodegeneration in a model of high adiposity with impaired glucose and insulin homeostasis. We compared this to mice on a nutritionally matched control diet (CD), and either the presence or absence of the metformin.

We found that metformin significantly delayed onset of motor and clinical signs and reduced the severity of disease signs at earlier time points. In HFD-fed mice, measures of both survival and health span were increased in metformin-treated mice relative to vehicle controls, as measured by motor signs of neurodegeneration. In CD-fed mice, metformin also showed a strong beneficial effect, improving health span. HFD, in the absence of metformin, was also mildly protective in terms of health span but did not extend survival. Since metformin is one of the most widely used drugs worldwide and is safe and cost-effective, our study suggests that it should be considered for repurposing for neuroprotection in both individuals suffering from obesity and diabetes, and in metabolically healthy individuals at risk of developing dementia later in life.

## RESULTS

### High-fat diet induced obesity is comparable to human adiposity with diabetic features by 24 weeks

Since mid-life obesity is associated with dementia later in life (Kivimäki *et al*., 2018; Floud *et al*., 2020), we first asked whether a mouse model of diet-induced obesity (DIO) could induce sufficient obesity and diabetic features to mimic the chronic exposure observed in people, but within the typical survival duration (approximately 28 weeks post-inoculation) of wild type C57BL/6J mice infected with RML scrapie. Un-inoculated C57BL/6J mice were thus fed a 60% HFD over 24 weeks, from 8 to 32-weeks of age (Figure 1A). These mice increased body weight by more than 75% after 20 weeks of HFD compared to mice fed with an ingredient-matched CD (Figure 1B). We specifically chose a CD to minimise differences to HFD, as most research to date has used chow diet with different mineral, vitamin, protein and fibre content and is a potential confounder. The weight distributions of CD- and HFD-fed mice were clearly separable at week 24 (Supplementary figure 1A). An average of 55% of weight gain was driven by increased fat mass, as determined by TD-NMR (Figure 1C and Supplementary Figure 1C). This increased adiposity was accompanied by impaired glucose tolerance (Figure 1D), impaired insulin tolerance (Figure 1E), and hyperinsulinemia (Figure 1F), as well as increased circulating triglycerides, cholesterol, LDL, and HDL (Supplementary figure 1D-F) after 20 weeks on the diet. Data showing no changes in lean mass between groups are shown in Supplementary figure 1G alongside example images of mice on CD or HFD, in Supplementary figure 1H. These features mimic the conditions of high BMI and adiposity and impaired glucose homeostasis seen in many humans with obesity.

**Figure 1.**
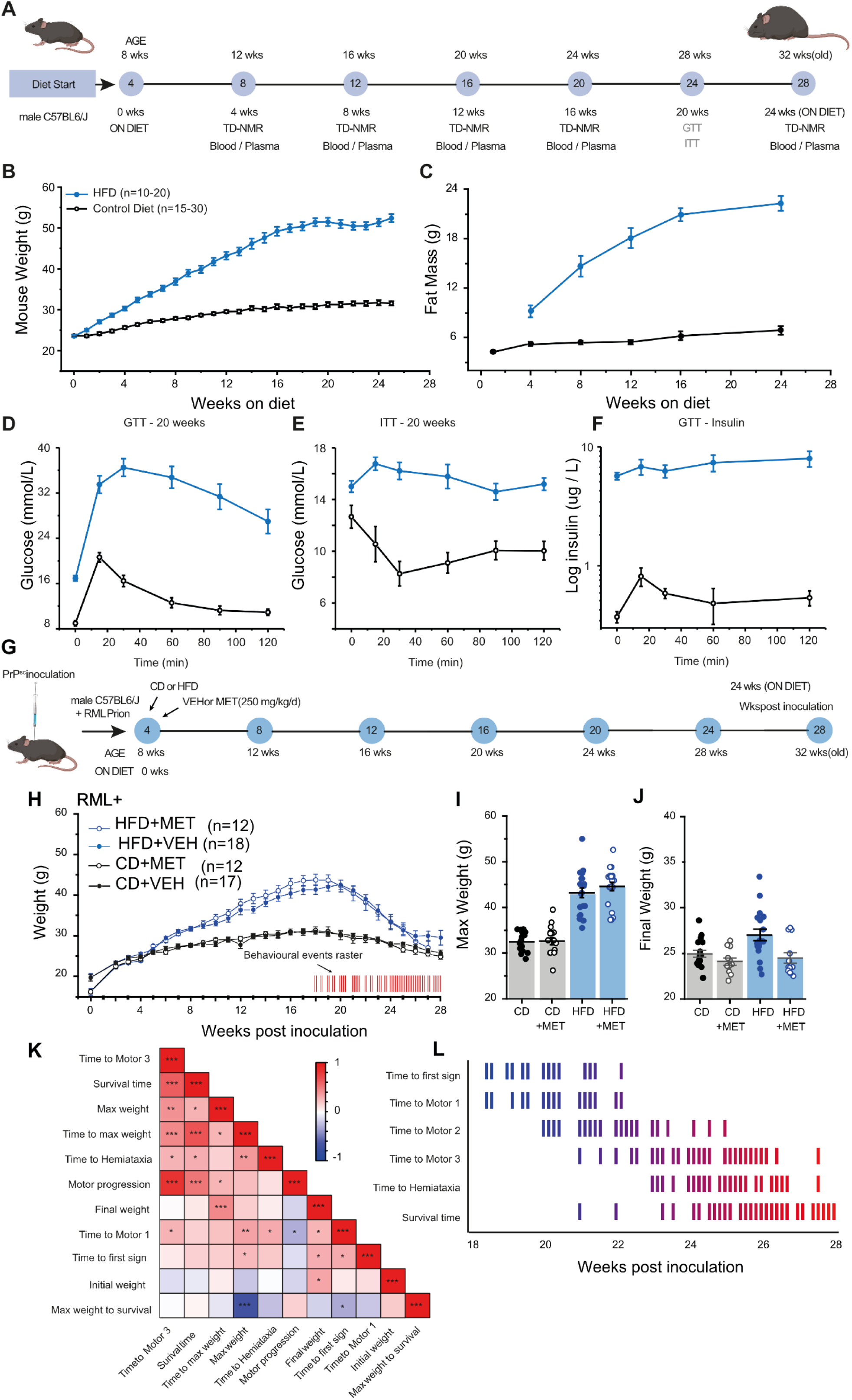
Metabolic parameters and neurodegenerative signs in HFD- and RML prion-treated mice. **A)** Experimental schematic of diet induced obesity (DIO) in male C57BL/6J mice. **B)** Increase in body weight of 60% in HFD-fed mice (n=10-20) compared to ingredient-matched control diet (n=15-30) starting at 8 weeks old. **C)** Accumulating adiposity in HFD-fed mice compared to CD-fed mice measured in fat mass (g) every 4 weeks. **D-F)** At 20 weeks, HFD-fed mice displayed impaired glucose tolerance (D), insulin tolerance (E) and hyperinsulinemia (F) relative to CD-fed mice. **G)** Experimental schematic by which RML scrapie-injected mice were randomly allocated into CD or HFD for 24 weeks and treated with vehicle or metformin (250 mg/kg/d) and maintained until developing confirmatory signs. **H)** Body weight measurements from groups described in (G). CD+VEH (n=17), CD+MET (n=12), HFD+VEH (n=18), HFD-MET (n=12). Inset raster plot (red) shows all behavioural events observed in inoculated mice aligned to the time course of body weight change. **I)** Distribution of body weights at wk 20, n numbers as per H. **J)** Distribution of body weights preceding confirmation, n numbers as per H. **K)** Pearson’s correlation coefficients of behavioural indicators plotted as a heatmap from -1 (Blue) to 1 (Red). Stars indicate statistical significance (0.05*, 0.01**, 0.001***). **L)** A raster plot of behavioural categories plotted over time showing progression of signs of neurodegenerative disease from 18 weeks post inoculation until all mice had reached confirmation. Colour gradient represents time scaled by the x axis. All graphs are plotted as mean ± SEM.

### RML scrapie-induced neurodegeneration interacts with HFD-induced obesity to induce rapid weight loss

Next, we inoculated male C57BL/6J mice with RML scrapie at four weeks of age and then four weeks later randomised them to HFD or CD treatment groups as per the schematic in Figure 1G. Since female mice in this model system frequently develop bladder enlargement resulting in death, we did not use mixed sex groups. One week after diet allocation, mice were further randomised to metformin (250 mg/kg/d) in the drinking water or a vehicle control (water). RML inoculated mice on HFD, both with and without metformin, rapidly gained body weight which peaked after 18 weeks post-inoculation (slightly earlier than non-inoculated mice in Figure 1H). From the non-inoculated experiments, we can deduce that the 40% increase in weight was primarily fat mass. Distribution of weights at week 20 are shown in Figure 1I. Following their attainment of peak weight HFD-fed mice, with and without metformin, underwent a rapid and sustained decline in body weight of approximately 0.35 g/day (0.88% of max/day). RML-inoculated mice on CD also peaked at the same 18-week time point (Figure 1H). Notably, CD-fed mice, with and without metformin, also declined in body weight at a lower rate of approximately 0.1 g/day (0.3% of max/day). Both groups appeared to reach a body weight plateau at around 26 weeks post-inoculation (30 weeks of age) likely due to the loss of adipose tissue. Distribution of weights at week 28 are shown in Figure 1J. In the last three weeks (26-28 weeks), all mice, regardless of disease severity, received additional wet-diet supplementation to mitigate any physical challenges associated with acquiring and consuming pelleted food. At the time that confirmatory signs of prion disease were observed, and animals were culled, HFD-fed mice were on average only 1.8 g heavier than CD-fed mice. HFD-fed mice were significantly (2.5 g) heavier than HFD-fed mice on metformin suggesting an interaction between diet and treatment at these late-stage time points (Figure 1J).

### RML-inoculated DIO mice demonstrate multiple behavioural indicators of neurodegeneration

Multiple methods have been used to classify the stage and time course of scrapie-induced neurodegenerative disease progression. Certain indicators facilitate end-point decisions (clinical signs) that are critical to experimental interpretation. Similarly, some indicators are secondary measures of disease severity. We set out to determine, in an unbiased manner, which indicators provided the most informative measurements of disease progression. To this end, we quantified a range of indicators (e.g. motor, behavioural, visual) and derivative metrics by frequent longitudinal sampling. We use ‘indicators’ to refer to any recorded observation of an RML-inoculated mouse at any stage or severity. These could then be systematically assessed which indicators could predict disease progression and survival. In all, 42 behavioural indicators and nine resulting metrics were assessed for each mouse at each observation attempt. These regular observations were performed from the development of the first sign to the time that clinical confirmatory signs of prion disease appeared over a time course of approximately 12 weeks. Overall, 60 RML-inoculated mice displayed 913 behavioural indications across approximately 3480 observations. These indications are summarised in the raster plot at the bottom of Figure 1H. A full list of clinical signs and other behavioural indicators is shown in Supplementary Figure 3.

We found that the start of behavioural indications coincided with the start of weight loss (event raster inset, Figure 1H). In addition, we found a clear significant and positive correlation between the appearance and progression of motor indicators (Motor progression, Motor 1 and 3, Hemiataxia and ‘time to maximum weight’) with later confirmatory signs (survival time) allowing us to identify those early signs that were most indicative of disease state. The Pearson’s correlation of indicators is plotted in Figure 1K. Importantly, some early signs such as loss of motor coordination have intrinsically high temporal variation until staged (M1, 2 or 3), at which point they show progressive loss of motor coordination that predicts survival (Figure 1L). Finally, we noted that impaired righting reflex, as used widely in the literature (Mallucci, 2009), was by far the most common confirmatory sign and a clear extension of progressive motor impairments. Conversely, a persistent hunched posture was a potentially false confirmatory sign since it appeared most frequently in mice with enlarged bladders observed on autopsy. Mice with enlarged bladders were excluded from the dataset.

### Metformin slows disease progression in RML-inoculated DIO mice

We initially hypothesised that HFD treatment alone would be sufficient to accelerate neurodegeneration compared to CD. We first analysed survival up to the point of robust confirmatory signs such as impaired righting-reflex, ataxia (sufficient to impair eating) and paw-dragging behaviour, but we did not observe a significant difference in survival between CD- and HFD-fed mice (Supplementary Figure 2A-E). Instead, the average time to first loss of motor coordination in HFD-fed mice was delayed relative to CD-fed mice, suggesting a protective effect restricted to early motor coordination (Supplementary Figure 2F). However, when we analysed whether metformin, given to mice on HFD, could alter neurological or behavioural outcomes we found that HFD-fed mice on metformin had significantly increased survival compared to HFD alone (Figure 2B) and also showed a significant delay in the mean time to confirmatory sign (Figure 2C). As expected, HFD-fed mice exhibited weight loss measured at their final weight, but it was also altered by drug treatment with reductions of - 37.2% for HFD-VEH and -44.7% for HFD-MET from maximum, such that metformin treated mice weighed more than those on vehicle (Figure 2D). Given the strong correlation between motor progression and survival, we selected two indicators of motor control to analyse disease progression prior to the appearance of confirmatory signs. We interpret these early and relatively minor motor signs as indicative of a better quality of life, so any delay in progression to more severe motor signs would be beneficial. We considered the first appearance of impaired motor coordination (M1) but saw no significant change by log rank test (Figure 2E) or in mean time (Figure 2F). We also considered the time to reach the final motor stage (M3) and found this was significantly extended in the HFD-MET group (Figure 2G). Time to the final motor stage was also significant by log rank test between these groups (Figure 2H). The mean distribution is shown in Figure 2I. Lastly, Given the overall correlation between survival and loss of final motor coordination, we considered whether this relationship changed when comparing treatment groups alone as shown in the scatter plot. We noted a clear change in the slope of the relationship while retaining the correlation (Figure 2J). Together, these data suggest that metformin can significantly extend survival and delay the progression of motor symptoms.

**Figure 2.**
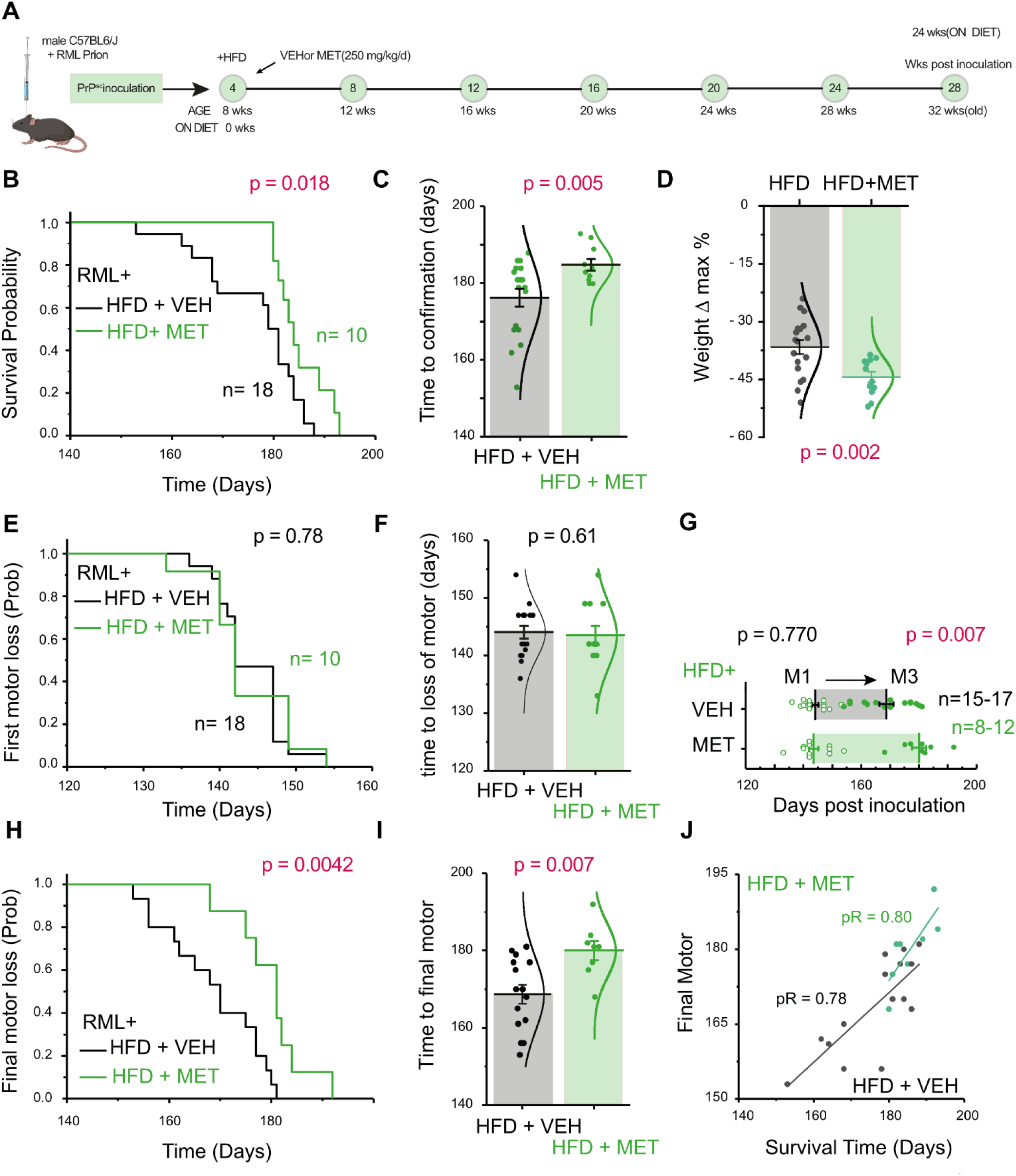
Survival and motor analysis of HFD-fed mice with and without metformin. **A)** Experimental schematic of inoculation with RML scrapie at four weeks, HFD treatment, random allocation vehicle or metformin, and behavioural monitoring until the development of confirmatory signs of prion disease. **B)** Kaplan-Meier curve of time to confirmatory sign for RML inoculated mice with CD+VEH vs CD+MET (χ2= 5.58 df = 1). **C)** Time to confirmatory sign of RML inoculated mice with CD+VEH vs CD+MET, (t = -3.07674 df = 25.75). **D)** Change in weight of RML inoculated mice with CD+VEH vs CD+MET (t = 3.425 df = 27.911). **E)** Kaplan-Meier curve of time to loss of motor coordination for RML inoculated mice with CD+VEH vs CD+ME, not significant. **F)** Time to loss of motor coordination for RML inoculated mice with CD+VEH vs CD+MET, not significant. **G)** Motor progression by severity stage (M1 to M3) for RML inoculated mice with CD+VEH vs CD+MET, Stage M1 n.s, Stage M3 (t =-2.968, df =21). **H**) Kaplan-Meier curve of time to final motor sign (M3) (χ2= 8.18, df = 1). **I)** Time to final motor sign (M3), (t = -2.968 df = 21). **J)** Scatter plot for survival time vs time to M3 for CD+VEH vs CD+MET, where pR is Pearson’s R. All graphs are plotted as mean ± SEM.

### Metformin slows disease progression in metabolically healthy RML-inoculated mice

As metformin was able to delay neurological and clinical outcomes across the time course of neurodegeneration and weight loss, we wondered to what extent it would be protective in metabolically healthy mice fed on CD (Figure 3A). These CD-fed mice with or without metformin did not show significant differences in survival by log rank (Figure 3B) but we did observe an increase in group mean survival time for the CD+MET group (Figure 3C). As expected, weight loss was a prominent feature of the neurodegenerative time course with CD and CD+MET groups losing 22.4% and 26.4% respectively body weight compared to the maximum weight they attained (Figure 3D). Next, we considered the time to first loss of motor coordination, as a strong correlation of disease progression. Here the CD+MET group showed significantly extended time to motor impairment compared to CD-fed mice by a log rank test (Figure 3E), as well as in the mean time for each group (Figure 3F). We also analysed motor progression across the CD and CD+MET groups, as we had done for mice fed HFD, by considering the time to appearance of impaired motor coordination (M1) and the final motor stage (M3) plotted in Figure 3G. We did not observe any differences between these groups. Time to the final motor stage was also significant by log rank test between these groups (Figure 3H). The mean distribution is shown in Figure 3I. Lastly, given the overall correlation between survival and loss of final motor coordination, we considered whether this relationship changed when comparing treatment groups alone. This is shown in the scatter plot (Figure 3J). Together, these results indicate that without the presence of obese or diabetic phenotypes, metformin was sufficient to slow the progression of motor signs of prion disease.

**Figure 3.**
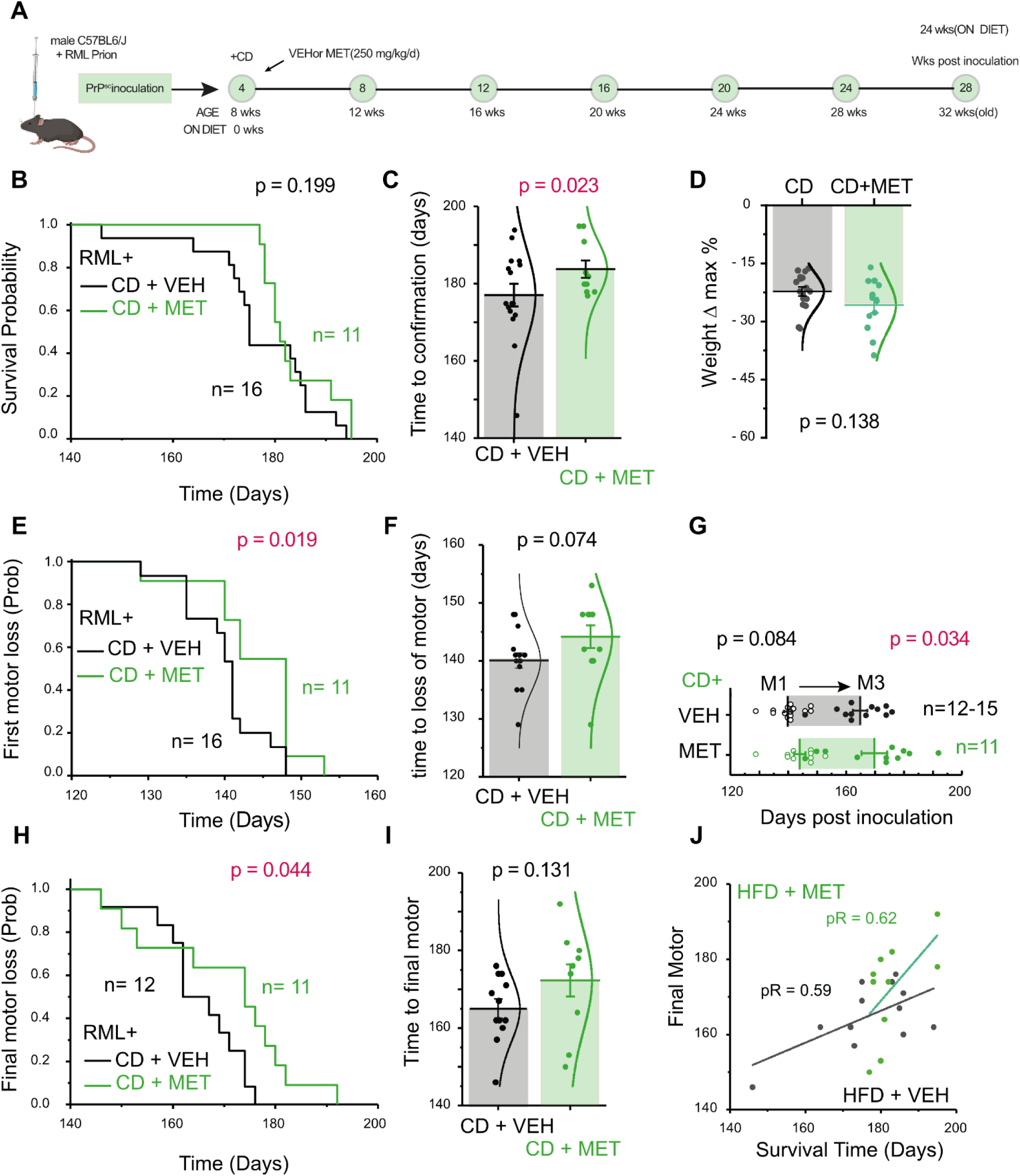
Survival and motor analysis of CD-fed mice with and without metformin. **A)** Schematic *RML-CD-VEH* and *RML-CD-MET* inoculated with RML scrapie at 4 wks and randomised *from* chow diet to CD, with or without metformin (MET) treatment (200 mg/kg/d by drinking water), at 8-wks old and maintained until developing confirmatory signs. **B)** Kaplan-Meier curve of time to confirmatory sign for RML inoculated mice with CD+VEH vs CD+MET (χ2= 2.375, df = 1). **C)** Time to confirmatory sign RML inoculated mice with CD+VEH vs CD+MET (t = 2.126, df = 23.85). **D)** Change in weight of RML inoculated mice with CD+VEH vs CD+MET (t = 1.6597 df = 27). **E)** Kaplan-Meier curve of time to loss of motor coordination for RML inoculated mice with CD+VEH vs CD+MET (χ2= 5.509, df = 1). **F)** Time to first loss of motor coordination for RML inoculated mice with CD+VEH vs CD+MET (t = -1.772, df = 17.66). **G)** Motor progression by severity stage (M1 to M3) for RML inoculated mice with CD+VEH vs CD+MET, M1 and M3 not significant. **H)** Kaplan-Meier curve of time to final motor sign (M3) (χ2= 4.046, df = 1). **I)** Time to final motor sign (M3), not significant. **J**) Scatter plot for survival time vs time to M3 for CD+VEH vs CD+MET. pR is Pearson’s R. All graphs are plotted as mean ± SEM. P values and n numbers are shown on the figures.

### Metformin reduces markers of hippocampal ER stress and neuroinflammation

To better understand the action of metformin in RML-inoculated brains, we culled mice treated with either metformin or vehicle at 20 weeks post-inoculation (Figure 4A), performed immunohistochemistry on hippocampal sections for markers of microglia (IBA1) and astrocytes (GFAP) (Figure 4B), and took high magnification images to assess microglial morphology (Figure 4C). We observed little change in the density of Iba1-positive cells (Figure 4D) but saw a significant decrease in GFAP immunoreactivity (Figure 4E), as well as a reduction in indicators of microglial activation as indicated by their decreased soma area (Figure 4F) and circularity (Figure 4G) in the presence of metformin. Furthermore, since neuronal ER stress is a common feature of neurodegenerative diseases including scrapie models, we assayed for pPERK expression (Figure 4H-I). We observed a significant (P=0.014) and substantial decrease in pPERK immunoreactivity in metformin treated animals (Figure 4G). Together, these histological features of reduced ER stress and reduced neuroinflammation are consistent with the delayed appearance of clinical signs and suggest that metformin is acting in the brain to exert its neuroprotective effects.

**Figure 4.**
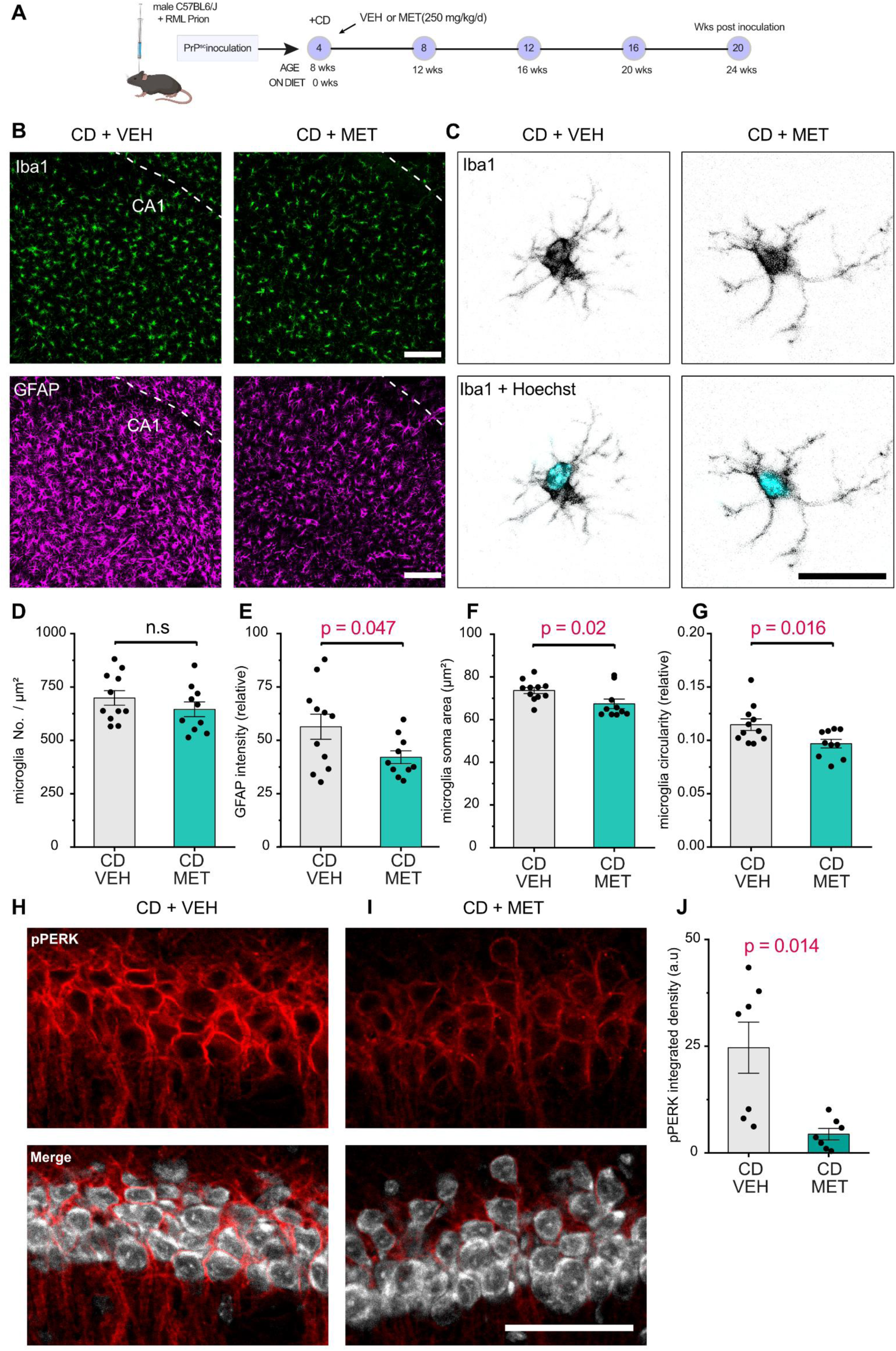
Metformin drives changes in microglia and reduces ER stress marker pPERK. **A)** Schematic for RML inoculation and metformin treatment over 20 weeks followed by immunohistochemistry on hippocampal sections. **B)** Examples of immunohistochemistry for microglia (Iba1) and astrocytes (GFAP) in CA1 between CD+VEH and CD-MET treated mice. **C)** Examples of microglia morphology changes between CD+VEH and CD-MET treated mice. **D)** Quantification of microglia numbers in CA1 per μm^2^ was not significant. **E)** Quantification of relative astrocyte numbers in CA1 using GFAP intensity (t = 2.162, df = 14.73). **F)** Quantification of microglia soma area (t = 2.387, df = 15.72). **G)** Quantification of microglia circularity (t = 2.59, df = 18.04). **H)** Example images of pPERK expression between CD+VEH and CD-MET treated mice (t = 2.162, df = 14.73). J) Quantification of pPERK in CA1 neurons. (t = 3,302, df = 6.62). All graphs are plotted as mean ± SEM. P values are shown on the figures. P values less than 0.0001 are shown as ****.

### Metformin penetrates deep structures in the mouse brain

The mechanism by which metformin slows prion disease progression is unclear. Since metformin had an apparently neuroprotective effect in both HFD-fed mice with impaired glucose homeostasis and in metabolically healthy CD-fed mice, we hypothesised that metformin might exert its effects directly in the brain rather than by normalising metabolic state. To play this role, metformin would have to cross the blood-brain-barrier (BBB) and accumulate in disease-relevant brain regions such as the hippocampus at sufficient concentrations. To determine if this could be the case we took in healthy (non scrapie-inoculated) C57Bl/6J mice and fed them CD or HFD with and without metformin for 20 weeks and tested for gene expression changes in the hippocampus (Figure 5A). We found that metformin modestly increased the expression of markers of astrocytes (Figure 5B) but not microglia (Figure 5C), but remarkably we observed changes in gene expression for ER stress markers including upregulated *Atf6* and *Chop* and downregulated *Xbp1* (Figure D-G). In HFD fed mice we also noted a decrease in *Rbm3* expression which was not seen when HFD fed mice that were treated with metformin. *Rbm3* expression is neuroprotective in scrapie inoculated mice, but its role here is unclear (Preußner *et al*., 2023). Overall, these findings corroborated our hypothesis that metformin can act directly in the deep brain, and on ER stress pathways.

**Figure 5.**
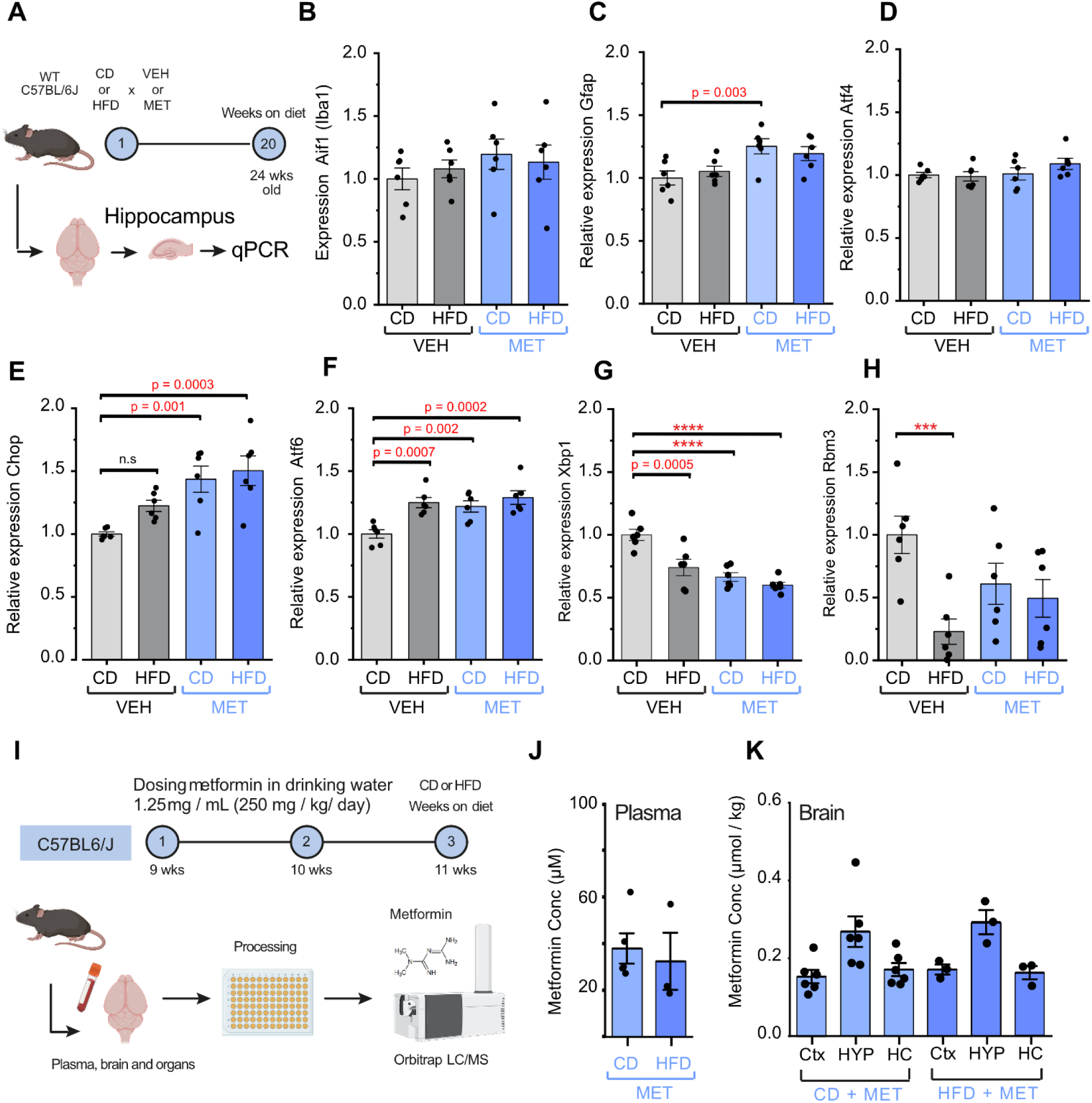
Gene expression in hippocampus of WT C57bl6J mice on CD or HFD and Veh or Metformin and concentrations of metformin in plasma, organs and brain. **A)** Schematic of WT C57bl6J mice on HFD or Metformin (250 mg/kg/d) for 20 weeks followed by qPCR on hippocampal tissue for relative gene expression of ER stress and neurodegeneration associated markers. **B)** Aif1. **C)** Gfap. **D)** Atf4. **E)** Chop. **F)** Atf6. **G)** Xbp1. **H)** Rbm3. Analysed by two-way ANOVA and bonholm post-hoc test. Plasma, brain and organ concentrations of metformin (250 mg/kg/d) after two-week administration in CD or HFD-fed mice**. I)** Schematic of metformin administration to mice for three weeks in drinking water (250 mg/kg/d before plasma and brain concentration measurements using CD-fed (n=6) and HFD-fed mice (n=3). **J)** Metformin concentrations in the plasma in µM. **(K)** Metformin concentration in the brain in µmol/kg. All graphs are plotted as mean ± SEM. P values and n numbers are shown on the figures. P values less than 0.0001 are shown as ****.

Since little seemed to be known about the biodistribution of metformin in specific brain structures such as the hippocampus, or how its distribution might be altered by the presence of excess adipose tissue, we then directly measured its concentrations in CD-fed and HFD-fed mice using targeted mass spectrometry. Specifically, we dosed CD- and HFD-fed mice with metformin at 1.23 mg/mL (∼250 mg/kg/d) in the drinking water as described above (Figure 5I) and measured metformin concentrations in the plasma, liver, enteric fat, muscle, and brain regions including the cortex, hypothalamus, and hippocampus after 3 weeks of treatment. Metformin concentrations in the plasma averaged 40 µM (Figure 5K), which is broadly equivalent to the metformin concentration in human plasma when given at common clinical doses (up to 2.5 g/d) (He and Wondisford, 2015). In the liver and muscle, we observed approximately 5 to 25-fold higher concentrations of metformin (∼1-3.5 µmol/kg) than in adipose tissue or the brain (∼0.2 µmol/kg), possibly by virtue of metformin’s hydrophilic properties. We did not observe any significant differences in metformin concentrations across diet groups. Measured metformin concentrations in the brain were highest in the hypothalamus (∼300 nmol/kg), and lower in the cortex and hippocampus (∼200 nmol/kg) (Figure 5L). Since metformin has been shown to inhibit cytokine release from vascular cells at concentrations as low as 1 nM (Isoda *et al*., 2006), and the concentrations we measured in the brain (∼200-300 nM) are substantially higher, these results suggest that it is plausible that metformin may exert its neuroprotective effects directly in the brain.

### Metformin does not cell-autonomously activate microglia at brain-relevant concentrations

Metformin may act in the deep brain and have secondary pharmacology separate from its action on systemic glucose control. However, the cellular basis for this interaction on microglia, astrocytes or directly on neurons is unclear. We hypothesised that since metformin changes morphological features of microglia in the complex *in vivo* environment, then a reduced and simplified *in vitro* system would shed light on whether metformin acts directly on microglia via cell-autonomous or non-cell-autonomous mechanisms. We therefore derived microglia from induced pluripotent stem cells (iPSC) and incubated them with vehicle or 250 nM metformin (Figure 6A) with or without the proinflammatory stimulus lipopolysaccharide (LPS), which is commonly used to acutely activate microglia. This concentration of metformin we used in this study was squarely within the range we measured directly in the mouse brain (200-300 nM). Using RT-qPCR, we found that in unstimulated microglia, metformin alone was not able to change expression levels of the genes *CD68* (Figure 6B), *IBA1* (Figure 6C) and *CD11B* (Figure 6D) or change the expression of markers of inflammation (Figure 6E-H). However, in the presence of LPS, metformin appeared to modulate microglial reactivity, suggesting that metformin at this concentration may act on microglia in an LPS-dependent or cell state-dependent mechanism (Figure 6E-H).

**Figure 6.**
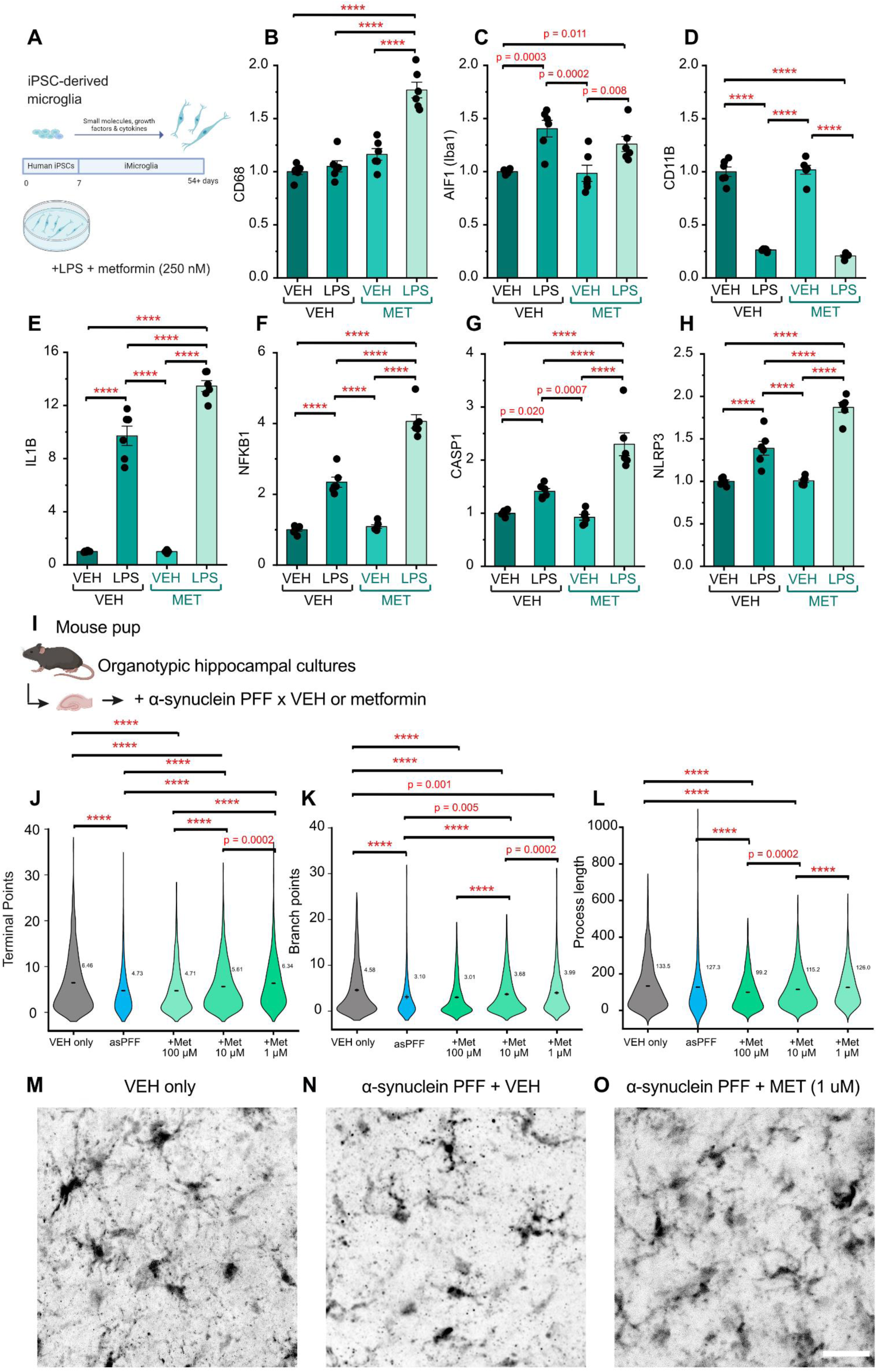
iPSC-derived microglia and mouse pup organotypic slices challenged with VEH or metformin in the presence of LPS or alpha-synuclein PFF respectively. **A)** iPSC-derived microglia incubated with VEH or MET (250 nM) in the presence or absence of LPS followed by qPCR microglia and inflammatory pathways. B) *CD68*. C) *AIF1*. D) *CD11B*. E) *IL1B*. F) *NFKB1*. G) *CASP1*. H) *NLRP3*. **I)** Schematic for an organotypic hippocampal slice culture system inoculated with a-synuclein PFF and treated with metformin at 1, 10 and 100 μM. **J-L**) microglial morphology parameters with and without a-synuclein PFF and treated with metformin at three concentrations including (**J**) No. of terminal points, (**K**) No. of branch points, (**L**) length of processes, **M-O**) Example images of microglial culture with and without a-synuclein PFF and the lowest concentrations of metformin (1 μM). All graphs are plotted as mean ± SEM. P values are shown on the figures. No p value means no significant difference. P values less than 0.0001 are shown as ****.

### Metformin acts directly on the brain to reduce neuroinflammatory markers in an alternative model of neurodegeneration

Since most of our studies were carried out in scrapie mice, it is difficult to clearly distinguish cell-autonomous or non-cell-autonomous mechanisms in the brain, from systemic drug mechanisms *in vivo*. To address this, we set out to test whether metformin could have neuroprotective effects in an alternative model of neurodegeneration in an *in vitro* culture system. Specifically, we selected a mouse hippocampal organotypic slice model that can be maintained for many weeks and can effectively model neurodegeneration when seeded with alpha-synuclein preformed fibrils (asPFF). This model system is known to cause both strong inflammation and progressive neuronal loss relative to control-treated slice cultures (Figure 6I) (Barth *et al*., 2021). To gain insight into the concentrations of metformin that might impact neurodegeneration-associated phenotypes, we incubated slice cultures (n=3 biological replicates/condition) not treated with asPFF as a negative control (VEH only), treated with asPFF alone (asPFF) or asPFF with three various concentrations of metformin (1, 10 and 100 μM) for 6 weeks. After this time, we fixed and immunostained slice cultures for microglia using IBA1 and quantified morphological features associated with inflammatory state. We found that metformin dose-dependently shifted microglial branching in the asPFF-seeding model. Surprisingly, we observed a dose dependent effect of metformin that was more beneficial at lower doses with microglial morphology better resembling the control conditions (Figure 6 J-L). Example images from each culture are shown in (Figure 6 M-P). These data suggest that lower doses of metformin, similar to those measured in the mouse brain are able to reduce cellular features of neurodegeneration, induced either by proteostatic stress or prion-like propagation of pathogenic alpha-synuclein. Together with our results from RML scrapie mice, these findings support the hypothesis that metformin can act directly in the deep brain to slow observed behavioural and molecular signs of neurodegeneration, without requiring systemic action on blood glucose control, via a mechanism that correlates with reduced markers of neuronal ER stress and neuroinflammation.

## DISCUSSION

Here, we show that metformin ameliorates the clinical signs of scrapie-induced neurodegeneration supporting the notion that it could be repurposed for neuroprotection in dementia. Importantly, we observed both prolonged survival as well as a reduced progression to early motor signs, suggesting a prolonged health-span. We do not find evidence that chronic adiposity and hyperglycaemia exacerbated disease progression in our model system (Østergaard *et al*., 2015; Zhu *et al*., 2022). Instead, the beneficial effects of metformin appeared to be either independent or potentiated by dietary status or metabolic health under the conditions tested. Diabetic patients treated with metformin have reduced risk of dementia (Samaras et al., 2020), and our findings that it is also neuroprotective in metabolically healthy mice suggests that its action on the brain may also confer benefit to a much broader population at risk of dementia. If true, the potential of repurposing this widely used and trusted compound beyond its typical treatment group would be substantial and could impact not only patients at risk of dementia, but also multiple neurodegenerative diseases. This type of broad scope therapeutic is urgently needed for the population-scale health problem of neurodegenerative disease, much as statins have revolutionised the management of cardiovascular disease.

Since metformin reaches deep brain structures at concentrations greater than required for activity *in vitro* (Isoda *et al*., 2006), and can be administered chronically over many months, we suggest that it may exert its effects directly in the brain, but further mechanistic studies are required to confirm our hypothesis. We speculate that several pathways may be plausibly involved, both directly and indirectly. First, we observe a reduction in the ER stress marker pPERK in CA1 neurons, a brain region commonly affected in human neurodegenerative diseases, at physiologically achievable concentrations. Metformin’s direct action on neuronal ER stress pathways may counteract disease-associated defects in proteostasis to prolong synapse integrity and neuronal health. We observed changes in these pathways in both the brain tissue of the scrapie model as well as in healthy mice, suggesting a direct action of metformin on these pathways that does not require underlying pathology. This contrasts with the action of metformin on human iPSC-derived microglia where metformin induced no perceivable response in the absence of LPS. And yet, in the scrapie model we observed changes in markers of neuroinflammation and microglial shape that we reproduced using an *ex vivo* model of a distinct neurodegenerative disease. It may be that metformin acts on microglia pathways in a pathology-dependent manner, but we are as yet unclear if these are upstream or downstream of changes in ER stress.

We also appreciate that metformin has many effects outside of the brain that may affect neurodegeneration. For example, it is an insulin-sensitising agent and there is evidence that insulin itself can restore hippocampal function to reverse cognitive deficits seen in obese mice (Gladding *et al*., 2018). Second, high levels of inflammatory TNFα are associated with dementia pathogenesis and insulin resistance in humans and are also necessary for insulin resistance in mice (Moller, 2000). This is also a feature of human Creutzfeldt–Jakob disease and mouse prion infection (Campbell *et al*., 1994; Sharief *et al*., 1999). High doses of metformin (3 g/d) in diabetic patients have been shown to reduce levels of circulating TNFα providing a potential indirect mechanism for metformin action in neurodegenerative disease. However, the effect of metformin on circulating pro-inflammatory cytokines is conflicting and with differing results at lower doses (<1.5 g/d) (Carlsen *et al*., 1998). In addition, TNFα^-/-^ mice are only resistant to prion infection when given non-cerebral inoculations, suggesting a possible mechanism specific to PrP^sc^ accumulation and not inflammatory pathways (Prinz *et al*., 2002; Aguzzi and Calella, 2009; Amoani *et al*., 2021). (Zhu *et al*., 2022). Lastly, circulating lipids in the plasma, particularly cholesterol, are a risk factor for dementia that are supported by evidence from Mendelian randomisation analysis (Østergaard *et al*., 2015; Zhu *et al*., 2022), and high levels of circulating lipids are reported in the CSF of people with AD (Byeon *et al*., 2021). Metformin can inhibit hepatic lipogenesis, but it is unclear whether it could function to address these risk factors specifically. The data we find here does not support an indirect mechanism given the absence of disease acceleration on HFD, despite high levels of circulating triglycerides and cholesterol, and the beneficial effects of metformin in organotypic slice cultures treated with asPFF.

Weight loss in people with dementia occurs both prior to diagnosis and in late-stage dementia and in approximately 40% of cases is a significant risk factor for death (Ciciliati *et al*., 2021). People with high BMI in mid-life can lose up to 5% body weight per year for the 8 years prior diagnosis with MCI or dementia (Franx *et al*., 2017; Singh-Manoux *et al*., 2018; Kang *et al*., 2021). The rate of body weight decline is also a predictor of later diagnosis. The association of weight loss and neurodegenerative disease creates an appearance of ‘reverse causation’ where high BMI in later-life is protective against dementia, whereas the opposite association is seen when considering BMI in mid-life (Li *et al*., 2022). RML-inoculated mice also display this disease-associated weight loss, which is most obvious in HFD-fed mice, and accompanies the earliest signs of disease (Figure 1H, J). Better understanding of the time course and mechanisms of weight loss in obese-RML models may shed light on the mechanisms behind neurodegeneration-associated weight loss in humans, as well as its value in predicting disease trajectory.

We found that mice fed HFD alone showed a small increase in the mean time to loss of motor control compared to CD-fed controls, but this was not observed in Kaplan–Meier survival analysis or other motor end points. In some mouse models with severe weight loss or wasting syndromes, such as the LMNA mutant Hutchinson-Gilford progeria line, lifespans are extended with an increased caloric intake (Kreienkamp *et al*., 2019). Similarly, in cases of human cachexia, increased calorie and/or protein intake is a recommended treatment (Gullett *et al*., 2011). However, in RML scrapie this is not as clear with HFD having only a modest effect on one measure of motor signs and not survival. Similarly, the combination of HFD and metformin caused a significant additive slowing of disease progression, while HFD and metformin fed mice weighed less, not more, than HFD-fed mice without metformin at the survival endpoint. We note that HFD in healthy mice, over 20 weeks, decreased the expression of the neuroprotective Rbm3 in the hippocampus (Preußner et al., 2023). Similarly, in the same mice, metformin prevented the HFD-induced decrease in Rbm3. But as we did not observe a worsening of motor endpoints with HFD further work is required to understand the role of Rmb3 in this context. Weight loss may occur through systemic inflammation via TNFα (Hennessy et al., 2017), or a result of overt neurological damage impairing the desire or ability to feed, but it remains poorly understood. One hypothesis is that metformin facilitates better usage of available fat stores through improved lipid trafficking and energy redistribution, in a manner that was not possible for HFD alone. More evidence will be required to confirm this.

We are unsure if there are sex-dependent effects in this model, since we had to exclude female mice, as many of them developed enlarged bladders in pilot studies. Small to moderately enlarged bladders in inoculated male mice seen on autopsy had no impact on outcomes across groups, but we believe that the detection and exclusion of mice with enlarged bladders is essential since the associated hunching behaviour can be mistaken as a confirmatory sign of prion disease. In the absence of enlarged bladders, hunching of any kind was rare in RML-inoculated mice C57BL/6J mice.

The identification of compounds that interact with known risk factors for dementia is an important potential treatment strategy. We note that newer treatments for obesity and diabetes, including GLP-1 agonists and semaglutide, that appear efficacious in retrospective analysis of randomised control trials for cardiovascular disease (Nørgaard *et al*., 2022), are now being advanced to phase 3 trials for AD in EVOKE (NCT04777396) and EVOKE Plus (NCT04777409) (Yun *et al*., 2018; Vargas-Soria *et al*., 2021). Metformin, as a potentially neuroprotective agent, will soon enter a multi-arm multi-stage trial for multiple sclerosis and maintains significant advantages in terms of cost and safety profile for dementia (Scherrer *et al*., 2019). In our view, dementia treatment at a population level will require broad scale public health intervention facilitated by safe and cost-effective medication such as metformin.

## MATERIALS AND METHODS

### Animals and colony maintenance

Animal work was performed under a Project Licence (PPL7597478) administered by the UK Home Office. All procedures were approved by the University of Cambridge Animal Welfare and Ethics Review Board (AWERB) and adhered to 3Rs and ARRIVE guidelines. All mice used in this study were of the C57BL/6J strain and were purchased at an age of four to six weeks from Charles Rivers Laboratories (Saffron Walden, UK). Upon arrival, mice were fed a standard chow diet and group housed in individually ventilated cages for seven days (non-inoculated in cages of five, RML and NBH inoculated housed in cages of three), to acclimatise them to the specific pathogen free facility. The animal facility was maintained on a standard 12-hour light/dark cycle (lights on at 0700h) and temperature was controlled at 22 ± 2°C. Water and food were available *ad libitum*.

### Diet and treatment group randomization and administration

All mice arriving remained on chow diet until age 7 weeks before acclimatisation to control diet (10% fat diet D12450Ji, Research Diets) for one week. Mice were then randomly allocated (8 weeks old) to either this control diet or a 60% high fat diet (D12492i, Research Diets). Diets were nutritionally balanced and matched to contain the same mineral mix, sucrose, protein and fibre content. Fat content was from the same source (predominantly lard), only in different proportions. Mice were weighed at least once weekly.

Prion-inoculated mice were also fed their allocated diets for approximately 24 weeks, until culled on presentation of terminal prion clinical signs (approximately 32± 2 weeks of age). Non-prion control mice undergoing metabolic characterization were maintained on diet for 24 weeks (until ∼32 weeks of age) before culling at a fixed timepoint.

At the end stages of disease, inoculated mice required wet-diet supplementation to facilitate feeding, without which they may die prematurely or exceed permitted severity. Dietary supplement was administered from week 20 post-inoculation and included high-calorie DietGel Boost (ClearH_2_0). Hydrogel or metformin-supplemented hydrogel was provided for hydration.

Metformin was added to the drinking water of appropriate cages at 1.25 mg/mL and replaced once weekly. This resulted in a dose of approximately 250 mg/kg/d metformin per mouse. The stability of metformin over the week and the administration dose was confirmed by LCMS (not shown). Control mice received normal drinking water using the same apparatus.

### Inoculation with brain homogenates

C57BL/6J male mice purchased at four weeks of age (Charles River) were acclimatised to the facility for one week in cages of three, and then randomly allocated for inoculation with either Rocky Mountain Laboratory (RML) prion or Normal Brain Homogenate (NBH). For inoculation, mice were anaesthetised with isoflurane and inoculated by injection in the parietal cortex with 30 µl of a 1% homogenate solution (gift from the laboratory of Prof. Giovanna Mallucci) as previously reported (Halliday *et al*., 2017). Mice were monitored until recovered and then returned to their home cage.

### Body composition (TD-NMR)

At four-weekly intervals, non-inoculated animals were assessed by TD-NMR. At this time tail-vein blood samples were collected for biochemical analysis using heparinised micro-haematocrit capillary tubes using standard protocols.

### Glucose and Insulin Tolerance Test (GTT/ITT)

For the glucose tolerance test mice (GTT) were fasted for 6 hours, and then given an intraperitoneal (IP) bolus of 20% glucose solution. Blood was sampled 30 mins prior to glucose injection (47829, Merck) followed by blood collection at 0, 15, 30, 60 and 120 minutes in heparinised micro-haematocrit capillary tubes (7493 11, Merck). These were centrifuged and the plasma flash frozen at -80°C and then stored at -20°C prior to submission for analysis. For the insulin tolerance test (ITT), performed at least one week after a GTT, mice were fasted for 6 hours prior to being given an IP bolus of 0.75 units/kg insulin (I9278, Merck). Blood was collected at 0, 15-, 30-, 60- and 120-minutes post-injection with heparinised micro-haematocrit capillary tubes. These were centrifuged and the plasma flash frozen at -80°C and then stored at -20°C prior to submission for analysis.

### Terminal tissue and blood collection

For all mice, a heparin-coated (375095, Sigma Aldrich) syringe with a 23G blunt-end needle was used to collect blood by cardiac puncture while under deep terminal anaesthesia, followed by cervical dislocation. Blood was transferred to tubes containing lithium heparin (450537, Greiner Bio-One MiniCollect™) and plasma was separated by centrifugation at 800 x g. Organs and brain tissues were immediately dissected and snap-frozen on dry ice.

### Clinical signs and observations

Mice were assessed for prion disease by the development of early and confirmatory prion signs as previously described (Moreno *et al*., 2013; Halliday *et al*., 2017; Smith *et al*., 2020). Mice were observed daily from the presentation of the first early sign to the appearance of confirmatory signs, and detailed assessments were made of all indications. Prion observations were blind to any previous records of early signs, but not blind to group variables. The presence of one early sign and a confirmatory clinical sign were sufficient to diagnose prion disease and terminate the mouse. All mice developed early signs prior to presentation of clinical signs. These are detailed in supplementary figure 3.

Inoculated animals are susceptible to bladder enlargement (small to large). On autopsy, all mice were checked for presence of gross pathology including blood, bladder enlargement or features of liver disease. ‘Small’-enlarged bladders were common in all groups, but urine could be manually expressed. On rare occasions ‘large’ enlarged bladders (detectable through the abdomen) presented as an un-sustained or sustained hunched posture, but with an inability to urinate. These were culled, confirmed on autopsy and excluded from survival data (<5% mice).

### Brain histology

RML scrapie-inoculated mouse brains were fixed in 4% PFA overnight and then decontaminated with >95% formic acid for 1 hour at RT. After refixation in PFA overnight, 25-µm thick brain sections were obtained using a Leica VT1000S vibratome (Leica Biosystems, Germany) within stereotaxic coordinates between 1.55 and 2.48 mm posterior to the bregma. Free-floating immunohistochemistry was conducted. For this, antigen retrieval in citrate buffer (pH=6) at +90°C (for pPERK staining) or autofluorescence quenching in 0.3% Sudan Black B in 70% ethanol for 20 minutes (for Iba1/GFAP staining) was performed. The sections were then treated with a blocking solution (10% NDS, Jackson ImmunoResearch Labs, USA; 1% BSA, Miltenyi Biotec Ltd, UK; 0.3% Triton X-100, Sigma-Aldrich, USA) for 1 hour at RT. Samples were incubated overnight at +4°C with primary antibodies: rabbit anti-Iba1 (1:750, Abcam, UK), chicken anti-GFAP (1:1500, Antibodies.com, UK), and rabbit anti-pPERK (1:250, Cell Signaling Technology, USA). After washing with PBS, sections were incubated with secondary antibodies for 1.5 hours at RT: donkey anti-rabbit AF488, goat anti-chicken AF647, and donkey anti-rabbit AF555 (1:1000; Invitrogen, USA). The brains were counterstained with NeuroTrace 640/660 (1:100; Thermo Fisher Scientific, USA) and Hoechst 33342 (1:10000; Invitrogen, USA), transferred onto histological slides, and mounted with Aqua/Poly-Mount medium (Polysciences, USA). Samples were visualised using a Leica SP8 confocal microscope (Leica Biosystems, Germany) at 20× and 63× magnifications.

### Orbitrap LC/MS for metformin

To measure metformin concentrations in mouse tissue (by weight) or plasma (by volume), tissue or plasma was extracted in 650 μL of chloroform and lysed at 6.5 m/s, 30 sec for 2 cycles (Tissuelyse). This was followed by 100 μL of metformin-d6 internal standard at 1 μM and 230 μL of methanol and vortexing, before adding 300 μL water. Samples were then centrifuged for 5 minutes at ∼20,000 *x g*. The aqueous fraction (top layer, 500 μL) was collected into a 2 mL amber-glass screw-cap vial before being dried down for 3 hrs at 60 °C and reconstituted in a low volume glass insert with 100 μL of 9:1 mobile phase in either 30 mM ammonium acetate in water with 0.02% acetic acid, or 20% acetonitrile and 80% water with 0.8% acetic acid.

Chromatography was performed using a Thermo Scientific Ultimate 3000 HPLC at a flow rate of 0.5 mL/min and Scherzo SM-C18 column (C18, cation & anion exchange: 150 mm * 3 mm I.D., 3 μm particle size) at 40 °C for 8 minutes and an injection volume of 5 μL. The needle wash was 20% acetonitrile, 80% water with 0.8% acetic acid and the gradient as follows: 0.0 minutes_10% [B], 0.2 minutes_10% [B], 1.2 minutes_99% [B], 5.0 minutes_99% [B], 5.1 minutes_10% [B], 8.0 minutes_10% [B].

To quantify the analyte, a 17-point metformin calibration curve using plasma as a matrix was prepared. The calibration level tested were: 10 nM, 15 nM, 20 nM, 25 nM, 50 nM, 75 nM, 100 nM, 200 nM, 500 nM, 1000 nM, 2000 nM, 5000 nM, 10000 nM, 20000 nM, 50000 nM, 75000 nM and 100000 nM. To evaluate the quality control of the curve, 8 metformin QCs were prepared using plasma as a matrix. The calibration levels tested were: 10 nM, 20 nM, 50 nM, 100 nM, 5000 nM, 25 μM, 80,000 μM.

The chromatography method was coupled to the MS tune page via Xcalibur software 4.1.31.9 and all the samples were analysed in a single sequence. The internal standard metformin-d6 was used to adjust the metformin quantification. For metformin standard (and experimental samples), mass range was 130.10872, retention time 2 min, smoothing points 9, Signal to noise (S/N) threshold 3.0, enable valley detection 5.0, minimum peak height (S/N) 3.0.

### Human iPSC-microglia

Human iPSC-microglia were generated based on the previously published protocol (Washer et al., 2022). The non-adherent microglial precursor cells were matured in maturation media (advanced DMEM-F12, Glutamax, penicillin-streptomycin, interleukin-34, granulocyte-macrophage colony-stimulating factor, transforming growth factor beta 1, and macrophage colony-stimulating factor) for 14 days prior to experimentation. Matured iPSC-microglia was treated with either metformin (250 nM) or Vehicle (DMSO) 1 hour prior to vehicle or LPS (100 ng/ml) challenge for 24 hours.

### RT-qPCR

Total RNA from treated iPSC-microglia was extracted using the RNeasy Plus Mini Kit (Qiagen) and half of the hippocampus from inoculated mice treated with vehicle or metformin was extracted using the RNeasy Plus Universal kit (Qiagen) according to the manufacturer’s instructions. Extracted total RNA (0.6-1 µg) was reverse transcribed into cDNA using the iScriptTM reverse transcription supermix. Target genes of interest were determined using the Taqman gene expression ‘assay-on-demandTM’ assays (Applied Biosystems, Foster City, CA) using FAM-labelled probes for target genes and *ACTB* (Hs01060665_g1) and *GAPDH* (Hs99999905_m1) as reference genes. The primer/probe targets were CD68 (Hs02836816_g1), *AIF1* (Hs01032552_m1), *ITGAM* (Hs00167304_m1), *IL1B* (Hs01555410_m1), *NFKB1* (Hs00765730_m1), *CASP1* (Hs00354836_m1), and *NLRP3* (Hs00918082_m1). For the mouse experiments, *Actb* (Mm02619580_g1) and *Gapdh* (Mm99999915_g1) were used as reference genes, and primer/probe targets were Aif1 (Mm00479862_g1), *Gfap* (Mm01253033_m1), *A*tf4 (Mm00515325_g1), Ddit3 (Mm01135937_g1), *Atf6* (Mm01295319_m1), *Xbp1* (Mm00457357_m1), *Rbm3* (Mm00812518_m1). The reactions were dispensed with Echo 525 Acoustic Liquid Handler (Beckman Coulter) and run on the QuantStudio™ 5 Real-Time PCR System in 384-well plates (Thermo Fisher Scientific). All expression assays were normalised to the geometric mean of the reference genes *GAPDH/Gapdh* and *ACTB/Actb* for human/mouse experiments, respectively.

### Hippocampal organotypic slice cultures

Hippocampal organotypic slice cultures were prepared using C57BL/6J (Jackson) WT mice according to previously published protocols (Daniel et al. 2005; Novotny et al. 2016). All animal experiments were performed in accordance with German Animal Protection Laws and were registered as N 04/19 M. Mouse pups, four to six days after birth, were decapitated and the brain transferred to a petri dish containing Minimum Essential Medium (MEM - Gibco) supplemented with 2 mM Glutamax (pH 7.35). The hippocampi were removed and cut into 350 µm thin slices using a tissue chopper (McIlwain). Intact slices were transferred onto sterile Millicell Cell Culture Inserts (Merck) (1 slice / insert) in 24-well plates (Corning) with 300 µl pre-warmed culture MEM (Gibco) + 20 % Horse Serum (Gibco), 1 mM Glutamax (Gibco), 0.00125 % Ascorbic Acid (Sigma-Aldrich), 0.001 mg/ml Insulin (Gibco), 1 mM CaCl^2^ (Sigma-Aldrich), 2 mM MgSO_4_ (Sigma-Aldrich), 13 mM D-Glucose (Roth), 100 U / ml Penicillin/Streptomycin (pH 7.28). The entire procedure was performed in a semi-sterile laminar flow hood with approximately 18 to 24 intact slices generated per pup. The cultures were kept at 37 °C and 5 % CO2 with full medium exchange three times per week.

a-synuclein pathology was induced in hippocampal organotypic slice cultures 10 days after preparation by a-synuclein pre-formed-fibrils (asPFF) (gifted by Ronald Melki) as described in Barth et al. 2021. In brief, 1 µl of 0.5 µg / µl asPFF was added onto each slice culture once and the cultivation was continued as usual. Metformin treatment was performed by adding the drug at three different concentrations (1, 10 and 100 µM) to the culture medium and started one day before the seeding with asPFF.

### Hippocampal organotypic slice cultures immunohistochemistry

To fix hippocampal organotypic slice cultures the culture medium was aspirated, slice cultures were washed once with PBS and then fixed with 300 µl ml of 4 % PFA each underneath and on top of the insert for 2 hours at room temperature. After fixation, PFA was removed and the slice cultures were washed three times for 15 min with PBS and subsequently stored at 4 °C in PBS for up to one month.

Slice cultures were carefully cut from the insert with a scalpel while they were still attached to the membrane in order to maintain top-down orientation for later imaging and stained in a 48-well plate (one separate well per condition). Slices were washed in PBS for ten minutes and transferred into 5 % Normal Donkey Serum + 0.3 % Triton X-100 in PBS. Slices were blocked for two hours at room temperature before incubation with the primary antibody overnight at 4 °C (goat-anti-Iba1, Novus Biologicals, 1:250) in 2 % Normal Donkey Serum + 0.3 % Triton X-100 in PBS. After incubation with the primary antibody, slices were washed three times in PBS for 15 minutes each before secondary antibody incubation. Slices were incubated with a secondary antibody in 1 % Normal Donkey Serum + 0.3 % Triton X-100 in PBS for two hours at room temperature and finally washed again three times in PBS for 15 minutes. Finally, slices were mounted onto TOMO adhesive glass slides by detaching them from the membrane with a brush, dried for 15 minutes at 37 °C and cover-slipped with Dako Fluorescence Mounting Medium. Slides were dried for 24h at RT before storage at 4°C.

### Data Analysis and microscopy

Data was analysed in either Origin Pro or R using custom scripts. Time to event data including survival was analysed using the Kaplan Meyer plot and log rank test. Other data was analysed using two-way t-tests of equal or unequal variance. Variance was determined by F test. For analysis of Pearson’s correlation an imputation procedure was performed using the MICE package in R with predictive mean matching (PMM) for 10 iterations of 10 imputations and known seed value. Imputation accounts for 43 values in 248 used for correlation analysis. Data without imputation values was then used for all proceeding survival analysis.

Mouse histology image analysis was conducted using Fiji ImageJ version 1.54f with custom macros to measure Iba1+ cell density (utilising the MaxEntropy Threshold), as well as the area and circularity of Iba1+ microglial somas. The “Integrated Density” function was employed to analyse GFAP+ and pPERK+ images following the application of a fixed threshold.

Hippocampal organotypic slice cultures imaging was performed using a Nikon Ti2-E+Yokogawa W1-SoRa-SpinningDisk confocal microscope with a x20 water-immersion objective (CFI APO LWD 20x WI LAMBDA S/0.95/0.95). Images were analysed with the ‘surfaces’ and ‘filaments’ tools of Imaris (v. 9.7.2; Bitplane). To reconstruct microglial processes, microglia were semi-automatically modelled using the ‘surfaces’ tool and all background signal outside of these surfaces was removed to facilitate subsequent reconstruction of microglial processes with the ‘filaments’ tool. Finally, morphological parameters of microglia were extracted from Imaris’ statistics function.

## Author Contributions

**E.C.H** designed and performed experiments, analysed data, produced visualisations, and drafted the manuscript. **H.-J. C.C** assisted in experimental design, performed experiments, assisted in manuscript preparation. **D.S^1,3^, M.W, V.M, A.S, C.R. S.A, D.S^4^** performed experiments. **D.K-V** supported experiments and assisted in experimental design. **F.T.M** conceived of the project, acquired funding and assisted in experimental design, data interpretation and manuscript preparation.

## Acknowledgments

We are grateful to Giovanna R. Mallucci for providing RML inoculum for experiments commenting on the manuscript. We appreciate the assistance in generating iPSC-derived microglia from Sam J Washer and Andrew R. Bassett. F.T.M. is a New York Stem Cell Foundation - Robertson Investigator (NYSCF-R-156) and is supported by the Wellcome Trust and Royal Society (211221/Z/18/Z) and a Ben Barres Early Career Acceleration Award from the Chan Zuckerberg Initiative’s Neurodegeneration Challenge Network (CZI NDCN 191942) which supported this work and financially supported E.C.H, H.-J. C.C., A.S., C. R., and S.A. We thank Paulina Guevara Dominguez and Albert Koulman and the Core Metabolomics and Lipidomics Laboratory (CMaLL) Facility for running the metformin mass spectrometry experiments, and the Disease Model Core (DMC) for assistance with mouse metabolic phenotyping. CMaLL and DMC are funded by the UK Medical Research Council (MRC) Metabolic Disease Unit (MRC_MC_UU_00014/5), a Wellcome Trust Major Award (208363/Z/17/Z), and a Wellcome DRP. For the purpose of open access, the authors have applied a CC-BY public copyright licence to any Author Accepted Manuscript version arising from this submission.

## SUPPLEMENTARY FIGURES

**Supplementary Figure 1.**
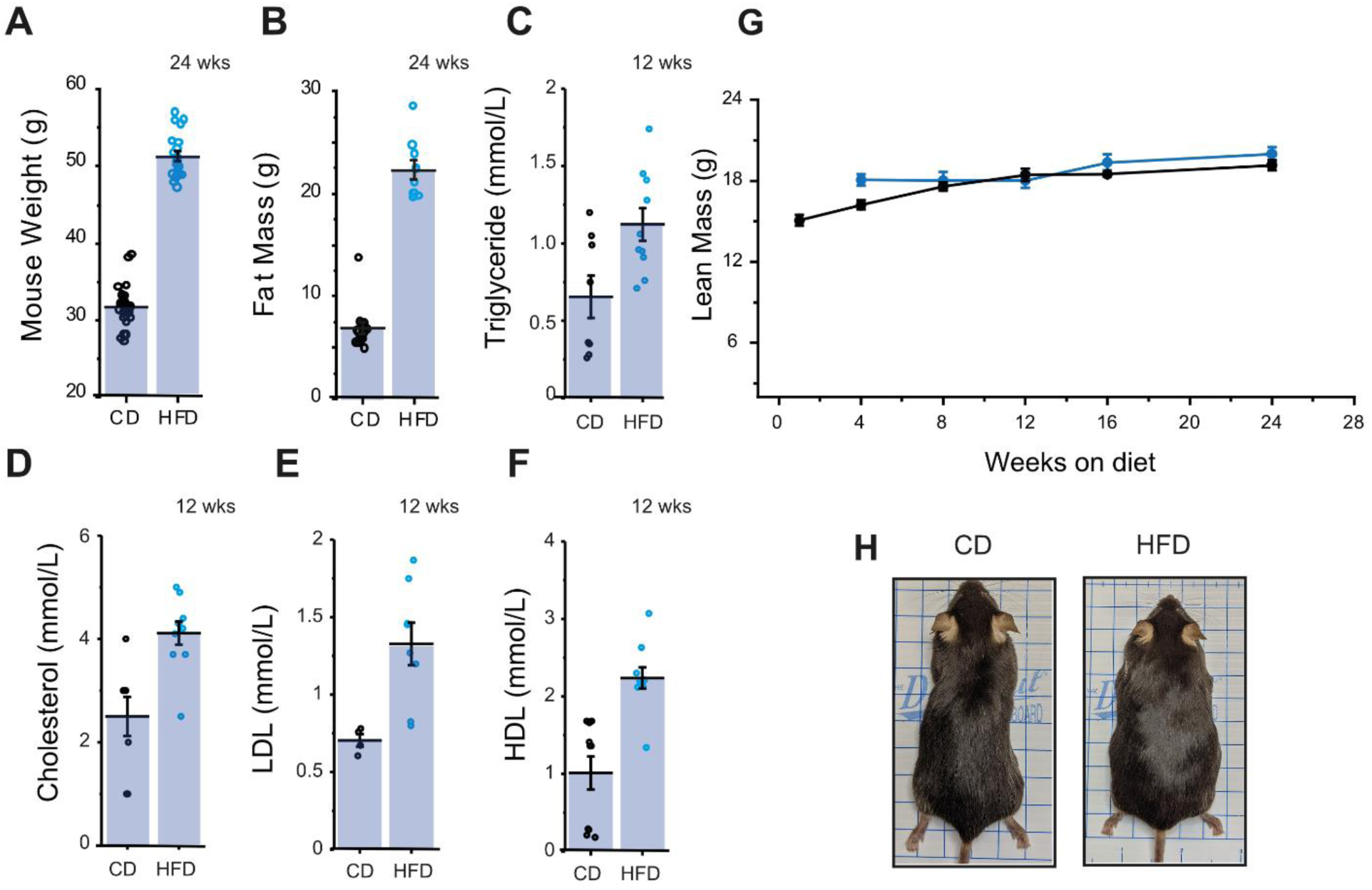
Effect of 24 weeks HFD in male C57BL/6/J mice on inducing obesity and metabolic syndrome supplementary data. **A)** Distribution of body weight at 24 weeks of 60% HFD fed mice (n=10-20) compared to ingredient-matched control diet (n=15-30) starting at 8 weeks old. **B)** Distribution of fat mass at 24 weeks of 60% HFD fed mice (n=10-20) compared to ingredient-matched control diet starting at 8 weeks old. **C)** Distribution of plasma triglycerides (mmol/L) at 24 weeks of 60% HFD fed mice compared to ingredient-matched control diet starting at 8 weeks old. **D)** Distribution of plasma cholesterol (mmol/L) at 24 weeks of 60% HFD fed mice compared to an ingredient-matched control diet starting at 8 weeks of age. **E)** Distribution of plasma LDL (mmol/L) at 24 weeks of 60% HFD fed mice compared to ingredient-matched control diet starting at 8 weeks old. **F)** Distribution of plasma HDL (mmol/L) at 24 weeks of 60% HFD fed mice compared to ingredient-matched control diet starting at 8 weeks old. **G)** Lean mass measured by TD-NMR over time. **H)** Example images of mice on CD or HFD. All graphs are plotted as mean ± SEM.

**Supplementary Figure 2.**
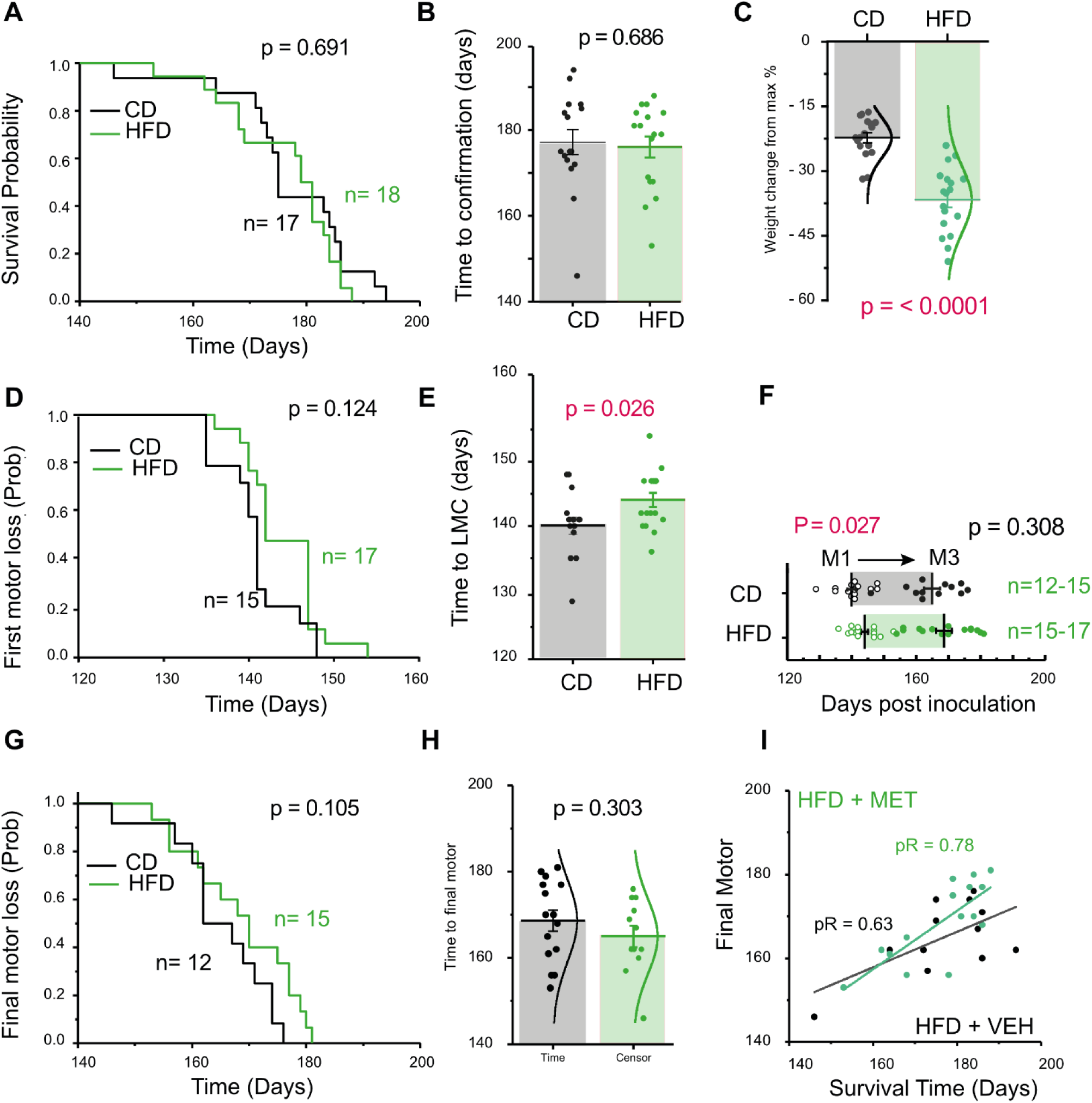
Survival and motor analysis of CD or HFD-fed mice. Experimental design is the same figure 2 and 3. **A)** Kaplan-Meier curve of time to confirmatory sign for RML inoculated mice with CD vs HFD was not significant by log rank test. **B)** Time to confirmatory sign in RML inoculated mice fed with CD vs HFD was not significant. **C)** Change in weight of RML inoculated mice fed with CD+VEH vs CD+MET was significant (t = 6.67 df = 33). **D)** Kaplan-Meier curve of time to loss of motor coordination for RML inoculated mice fed with CD+VEH vs CD+MET was not significant. **E)** Time to loss of motor coordination for RML inoculated mice fed with CD+VEH vs CD+MET was significant (t = -2.34 df = 30). **F)** Motor progression by severity stage (M1 to M3) for RML inoculated mice fed with CD+VEH vs CD+MET, is significant to M1 but not M3 (t = -2.34 df = 30). **G)** Kaplan-Meier curve of time to final motor sign (M3) was not significant). **H)** Time to final motor sign (M3), was not significant. I) Scatter plot for survival time vs time to M3 for CD+VEH vs HFD+MET. pR is Pearson’s R. All graphs are plotted as mean ± SEM.

**Supplementary Figure 3.**
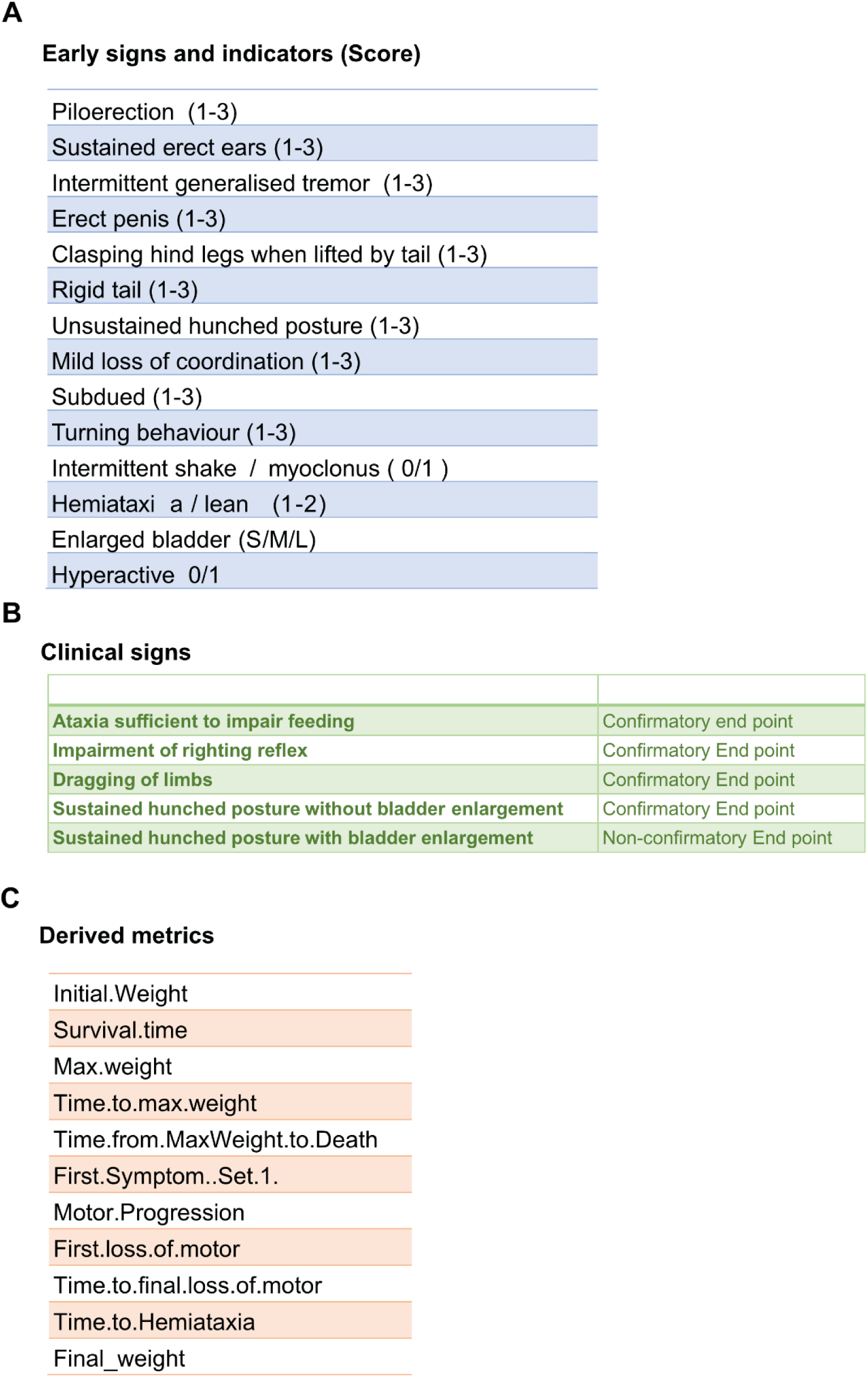
List of indicators used in the assessment of prion disease progression. **A)** List prion early indicators and early signs. (B) List of prion clinical signs. (C) List of derived metrics from prion early indicators and bodyweight data (C). Brackets indicate scoring per indication and associated outcomes are listed for each.

**Supplementary Figure 4.**
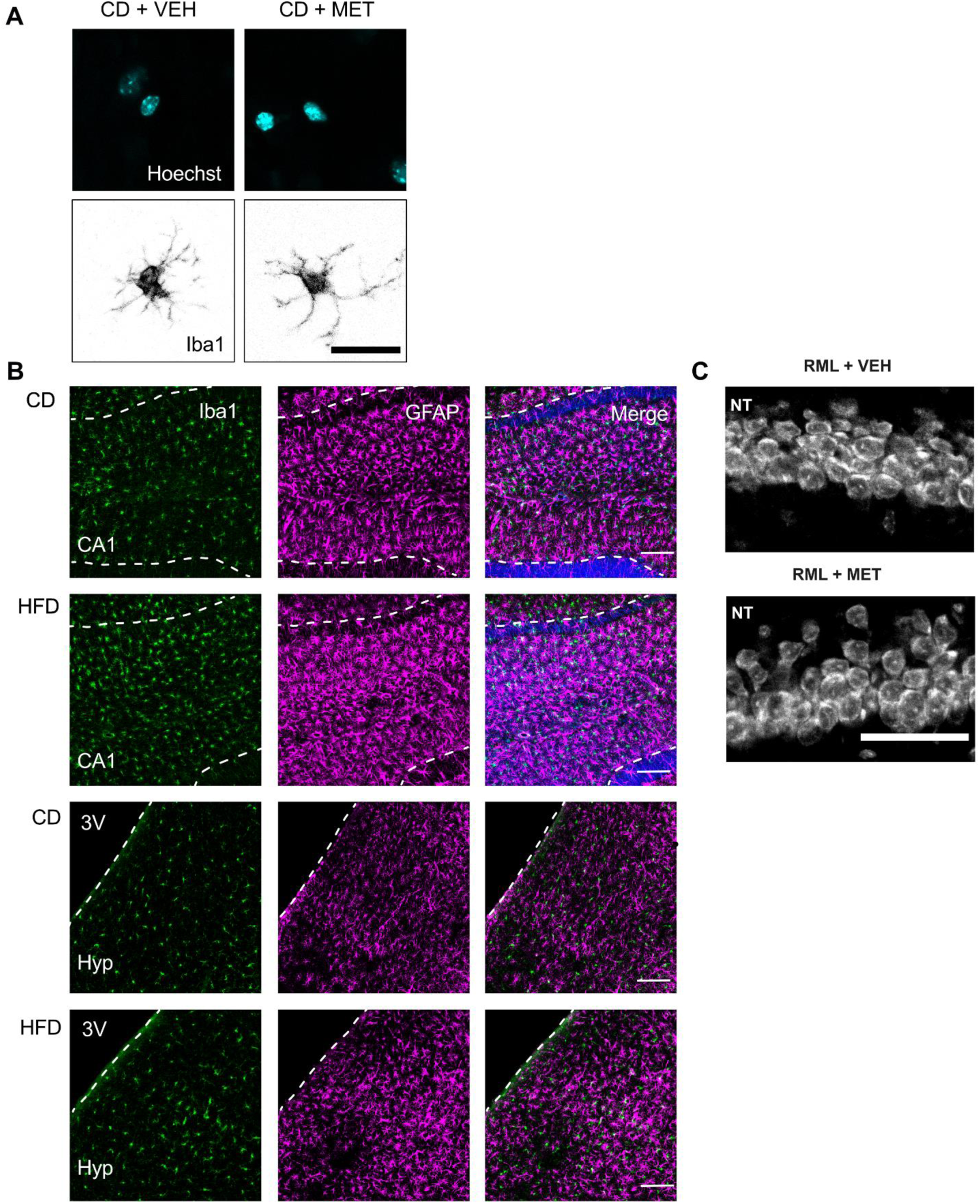
Additional images from high magnification panels and histology from CD vs HFD. A) Examples of microglia histology from Figure 4 showing separate channel hoechst with all nuclei B) Examples of microglia (Iba1) and astrocyte (GFAP) staining in CD vs HFD conditions in NBH mice after 20 on diet, across hippocampus (CA1) and the hypothalamus (Hyp). C) Example images of NeuroTrace (NT) from CA1 neurons used for pPERK example in Figure 4.

**Supplementary Figure 5.**
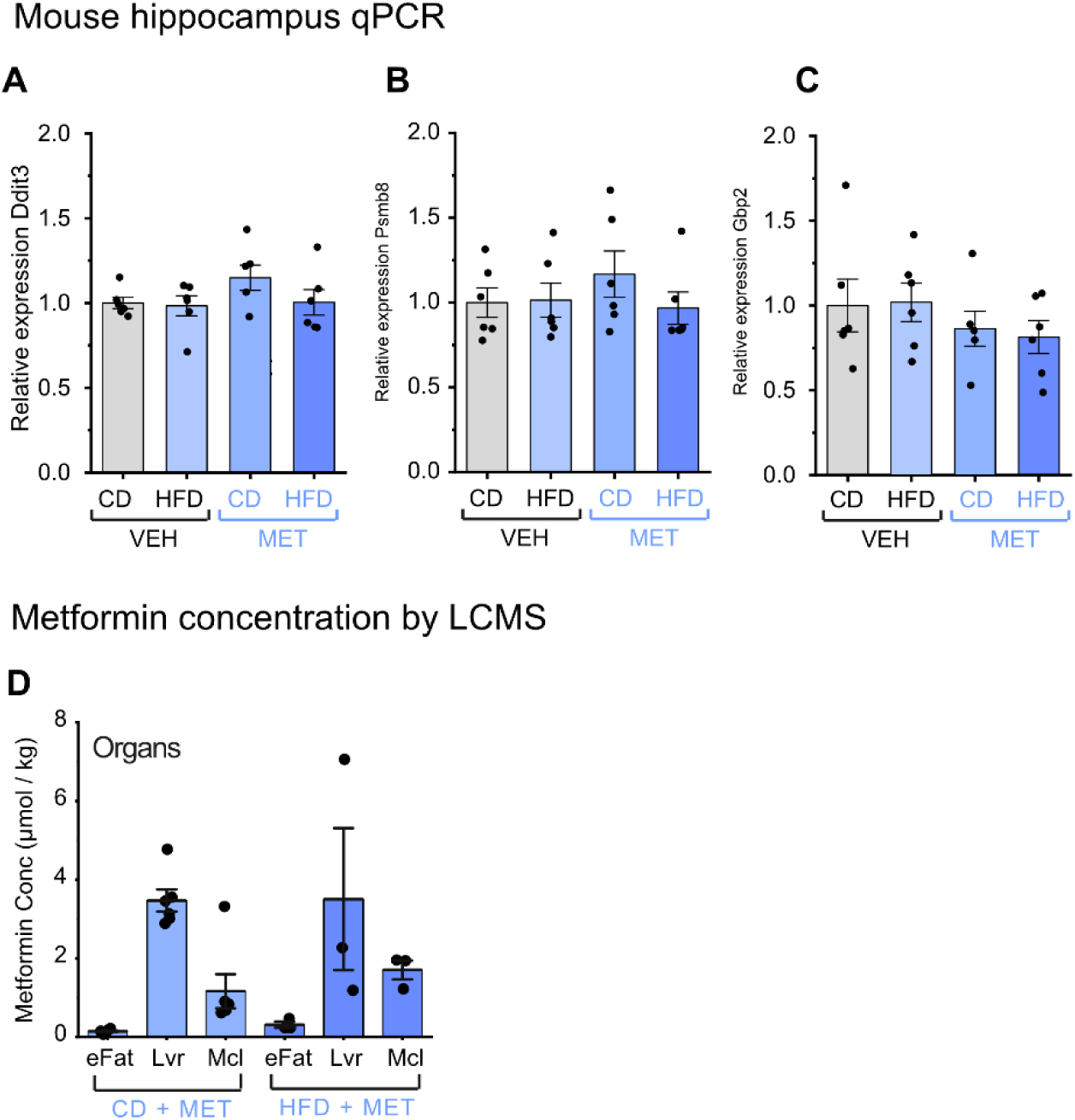
Relative gene expression by qPCR of WT mouse hippocampus after 24 weeks diet or drug treatment for: A) *Ddit3*. B) *Psmb3* C) *Gbp2*. **Metformin concentration by LCMS.** D) Metformin concentration in organs in µmol/kg including liver (Lvr), muscle (Mcl) and inguinal white adipose tissue (eFat) in µg/kg (L) in CD-fed (n=6) and HFD-fed mice (n=3). All graphs are plotted as mean ± SEM. P values are shown on the figures. Observational data on organs are not tested. P values less than 0.0001 are shown as ****.

**Supplementary Figure 6.**
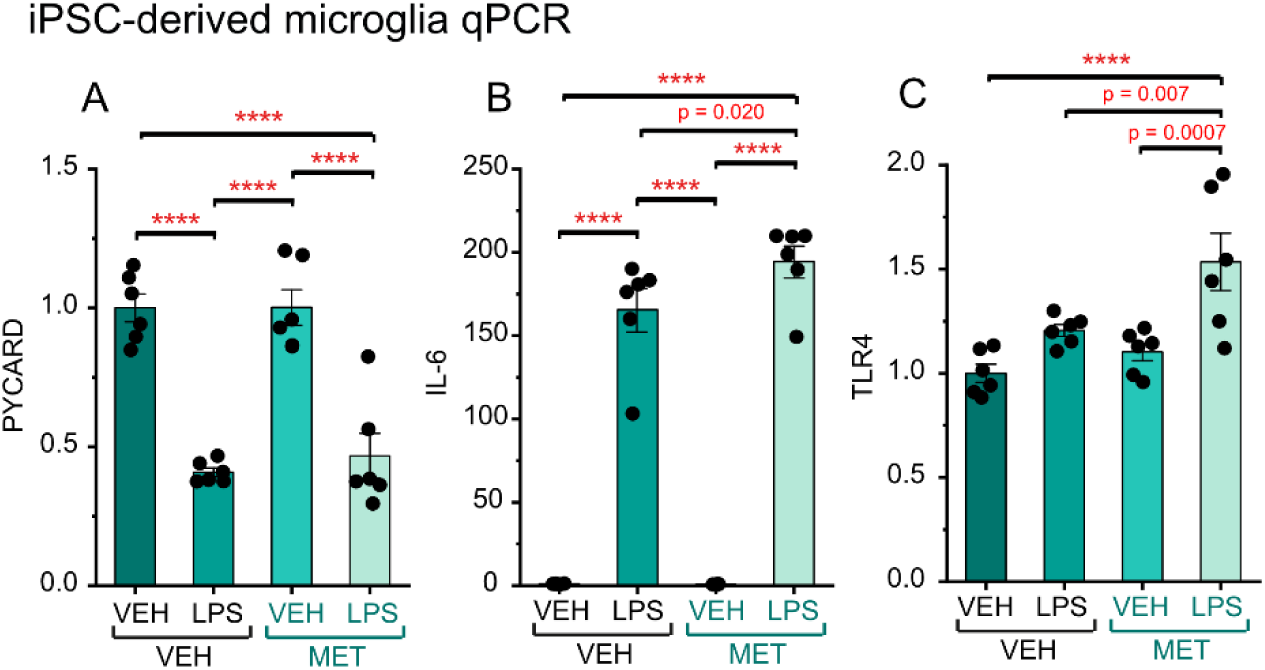
A) iPSC-derived microglia incubated with VEH or MET (250 nM) in the presence or absence of LPS followed by qPCR microglia and inflammatory pathways. A) *PYCARD*. B) *IL-6*. C) *TLR4*.All graphs are plotted as mean ± SEM. P values are shown on the figures. P values less than 0.0001 are shown as ****.

